# Strain and temperature dependent aggregation of *Candida auris* is attenuated by inhibition of surface amyloid proteins

**DOI:** 10.1101/2023.05.09.540062

**Authors:** Dhara Malavia-Jones, Rhys A. Farrer, Mark H.T. Stappers, Matt B. Edmondson, Andrew M. Borman, Elizabeth M. Johnson, Peter N. Lipke, Neil A.R. Gow

## Abstract

*Candida auris* is a multi-drug resistant human fungal pathogen that has become a global threat to human health due to its drug resistant phenotype, persistence in the hospital environment and propensity for patient to patient spread. Isolates display variable aggregation that may affect the relative virulence of strains. Therefore, dissection of this phenotype has gained substantial interest in recent years. We studied eight clinical isolates from four different clades (I-IV); four of which had a strongly aggregating phenotype and four of which did not. Genome analysis identified polymorphisms associated with loss of cell surface proteins were enriched in weakly-aggregating strains. Additionally, we identified down-regulation of chitin synthase and chitinase genes involved in the synthesis and dissolution of the chitinous septum. Characterisation of the cells revealed no ultrastructural defects in cytokinesis or cell separation in aggregating isolates. Strongly and weakly aggregating strains did not differ in net surface charge or in cell surface hydrophobicity. The capacity for aggregation and for adhesion to polystyrene microspheres were also not correlated. However, aggregation and extracellular matrix formation were all increased at higher growth temperatures, and treatment with the amyloid protein inhibitor Thioflavin-T markedly attenuated aggregation. Genome analysis further indicated strain specific differences in the genome content of GPI-anchored proteins including those encoding genes with the potential to form amyloid proteins. Collectively our data suggests that aggregation is a complex strain and temperature dependent phenomenon that may be linked in part to the ability to form extracellular matrix and cell surface amyloids.

**HIGHLIGHTS:** The amyloid inhibitor Thioflavin-T inhibited *C. auris* aggregation. Aggregating isolates do not exhibit any defects in cell separation.

Genomic differences were identified between strongly aggregating and weakly-aggregating strains of *C. auris*.

Aggregation did not correlate with surface charge or hydrophobicity of yeast cells.

## 1 Introduction

*Candida auris* has recently emerged as a global fungal pathogen whose antifungal drug and environmental stress resistant properties has resulted in its classification as a critical priority pathogen by the World Health Organisation (WHO 2022). Five genetically distinct clades of *C. auris* have been identified, which are somewhat geographically demarcated, although they are increasingly becoming globally dispersed (Arastehfar et al. 2021; Chow et al. 2019; Chow et al. 2020; Forsberg et al. 2019; Lockhart et al. 2017; Rhodes and Fisher 2019; Satoh et al. 2009). Several studies have revealed important phenotypic characteristics that may relate to the emergence of *C. auris* as an organism of major health care concern. In this study, we focus on the ability of some strains to form clumps of aggregated yeast cells.

Like many other *Candida* species, *C. auris* is a polymorphic fungus that is capable of limited filamentation (Kumamoto and Vinces 2005; Yue et al. 2018). In addition, some isolates in each of the major clades also exhibit the phenomenon of forming distinct cellular aggregates of yeast cells (Bing et al. 2023; Borman, Szekely, and Johnson 2016; Forgacs et al. 2020; Santana and O’Meara 2021; Szekely, Borman, and Johnson 2019; Zamith-Miranda et al. 2021). Recent studies have attempted to dissect the mechanism of aggregation and these have led to the identification of potential candidate genes that drive cellular aggregation (Bing et al. 2023; Santana and O’Meara 2021; Brown et al. 2020). Aggregating strains are more resistant to disinfectants, azole antifungals and to destruction by immune phagocytes and they have the ability to colonise and persist on biotic and abiotic surfaces. Furthermore, aggregating strains exhibit additional phenotypic differences with weakly-aggregating strains such as their capacity to form biofilms (Bing et al. 2023). Strongly aggregating strains are differentially recognised by immune cells and induce different host responses (Brown et al. 2020; Chakrabarti and Sood 2021; Sherry et al. 2017; Short et al. 2019) suggesting they may have different surface chemistries. While aggregating strains may be more difficult for immune phagocytes to destroy, they may also be compromised in their ability to disperse in the bloodstream. Virulence studies using the wax moth *Galleria mellonella* have demonstrated that aggregating strains are attenuated in virulence as compared to weakly-aggregating strains (Borman, Szekely, and Johnson 2016). This difference in virulence was also borne out by a later study in a neutropenic murine model, although large aggregations of cells were found in hearts, kidneys and liver of all mice suggesting that most strain will form aggregates in tissue (Forgacs et al. 2020). Aggregating strains also have been shown to cause an increased pro-inflammatory response on skin and at sites of wounds (Brown et al. 2020). These studies provide evidence of the clinical importance of aggregation in *C. auris* and the need to further dissect its aetiology.

Borman and colleagues reported that some *C. auris* daughter cells from aggregating isolates may not be released normally from mother cells post cytokinesis (Borman, Szekely, and Johnson 2016). It was later observed that deletion of ACE2 transcription factor in a weakly-aggregating *C. auris* strain was associated with decreased expression of chitinase gene *CTS1* that might participate in septal plate separation and hence aggregation (Santana and O’Meara 2021). Deletion of *ACE2* in *Candida glabrata* also resulted in clumping of cells and in this fungus the aggregating mutant was hyper virulent (Kamran et al. 2004). Therefore, aggregation could result from defects in cell separation.

Clade II strains of *C. auris* are mostly weakly-aggregating and lack fourteen adhesins found in other clades that play a role in adhesion to host surfaces (Chakrabarti and Sood 2021; Munoz et al. 2021). More recently, Als4 dependent aggregation was identified in some but not all *C. auris* aggregating isolates (Bing et al. 2023). It is therefore likely that aggregation is in part the result of the presence of specific surface proteins that play a role in adhesion. However, the relationship between adhesion and aggregation remains to be established and it is still not clear whether aggregation is caused by a combination of cell-separation defects or hyper-adherent properties of some *C. auris* isolates mediated by specific adhesins.

Some cell wall adhesin proteins such as Als5, Eap1, Hwp2 and Sap65 can form fibrous amyloid-aggregates by spontaneous conversion to a highly ordered β-aggregation state. The β-aggregation state of these cell wall proteins results is dramatic changes in their properties, including changes in their birefringence, ability to bind zinc, ability to promote biofilm formation and ability to undergo strong aggregation and the formation of cell-to-cell bonds (Kumar et al. 2017; Lipke et al. 2018; Lipke et al. 2021). The amyloid generation capacity of proteins of *C. albicans* is related to a seven amino acid sequence (IVIVATT) in the T-domain that can be used to undertake bioinformatic screens of genes with potential amyloid forming capacity.

Although the property of aggregation has been recently linked to ALS proteins (Bing et al. 2023), aggregation is not exclusively associated with particular *C. auris* clades, although *in vitro* it is most prominent in African clade III isolates. *C. auris* isolates belong to four common and one rare genetically distinct clade, which were each identified in different geographical locations. These clades are Clade I (South Asia), Clade II (East Asia), Clade III (Africa) and Clade IV (South America). The fifth clade was identified in Iran is separated from other clades by more than 200,000 SNPs (Chow et al. 2019). The four of five *C. auris* clades share an average pairwise nucleotide identity (ANI) of 98.7% (Munoz et al. 2018). However, Clade II genomes exhibit a high degree of rearrangements compared with clades I, III and IV including two inversions and nine translocations resulting in a substantially different karyotype (Chybowska, Childers, and Farrer 2020). Although the clades share a high ANI value, they are distantly evolved. Such genetic variations between the clades can have a significant impact on gene function resulting in a range of phenotypic variations, potentially including aggregation (Brown et al. 2020; Chybowska, Childers, and Farrer 2020; Lockhart et al. 2017). Whether the ability to aggregate has evolved once or multiple times in the *C. auris* species is yet to be determined.

In this study, we investigated the heterogeneity of aggregation in *C. auris* by comparing genomes and phenotypes of four strongly-aggregating and four weakly-aggregating *C. auris* clinical isolates belonging to the four different clades. Using whole genome sequences, biochemical, physiological and immunological analyses combined with live cell imaging, and scanning electron microscopy, we demonstrated that aggregating cells apparently undergo normal cell division although some genes involved with the formation and dissolution of the chitinous septum were down-regulated. The strongly aggregating and weakly-aggregating strains had similar cell surface charge, hydrophobicity, and adhesion properties and inhibition of amyloid proteins attenuated aggregation.

## 2 Materials and methods

### 2.1. Microbial strains and growth

*C. auris* clinical isolates were provided Prof. Elizabeth Johnson and Prof. Andy Borman, from the UK Health Security Agency. The isolates used and their associated clades are listed in Table 1. All isolates were stored as glycerol stocks at -80°C prior to use and were grown on Yeast-Peptone-Dextrose (YPD) agar for 24-48 h at 30°C and then stored at room temperature (RT) prior to propagation in YPD broth for 24 h at 200rpm, 30°C or 37°C. Following growth, all isolates were washed twice in sterile Millipore water. Cells were standardised to desired concentration using optical density measurements at OD_600_.

**Table 1.** Aggregation capacity and clade identity of clinical isolates used in this study.

| Isolate ID | Origin/Clade | Aggregation |
| --- | --- | --- |
| NCPF 8976 | South Asian (Clade I) | Weakly aggregating |
| NCPF 8977 | South African (Clade III) | Strongly- aggregating |
| NCPF 8983 | South Asian (Clade I) | Weakly aggregating |
| NCPF 8985 | South Asian (Clade I) | Weakly aggregating |
| NCPF 8996 | South African (Clade III) | Strongly- aggregating |
| NCPF 13029 | Japanese (Clade II) | Weakly aggregating |
| NCPF 13052 | South American (Panama) (Clade IV) | Strongly-aggregating |
| NCPF 13059 | South American (Colombia) (Clade IV) | Strongly-aggregating |

*C. auris* isolates reported to exhibit an aggregating phenotype were confirmed by suspending single colonies in 500 µl sterile Millipore water and imaged using DeltaVision microscope at 200× and 600× total magnification. To study effects of Thioflavin-T on aggregation, *C. auris* isolates were grown in the presence of 30 µM Thioflavin-T (dissolved in 10 mM Tris-EDTA, pH 7) for 24 h at 35°C in YPD. For experiments investigating the effects of temperature cells were grown in YPD at 30°C or at 37°C.

### 2.2. Microfluidics assay to study cell separation

All isolates were grown in liquid YPD at 30°C, 200 rpm for 16 h. Cells were washed twice with water and cell density was adjusted to an OD_600_ of 0.2 in 1 ml Millipore water. Then, 75 µl samples of culture were transferred to independent washed flow-inlet chambers of CellASIC ONIX Microfluidic plates. Additionally, 250 µl YPD broth containing 1 µg/ml Calcofluor White was added to the washed solution-inlet chambers to label the time=zero cell wall surface. Plates were sealed with CellASIC ONIX2 system manifold and prepared for live cell imaging. The CellASIC ONIX2 microfluidic system used an initial flow rate of 6 µl/sec. The pressure was set at 2 psi throughout the experiment to provide optimum media flow through the imaging microfluidic chambers. Using DeltaVision Elite at 100× total magnification, time lapse images were taken every minute for 8 h. The microscopy chamber was maintained at 30°C throughout the experiment which prevented filamentation. Following 8 h of incubation, time lapse images were analysed using ImageJ Fiji.

### 2.3. Alcian Blue assay for cell surface charge

Cell surface charge for the eight *C. auris* isolates was determined using the cationic Alcian Blue dye as previously described (Hobson et al. 2004). Briefly, overnight grown cells were harvested and washed twice with sterile PBS. They were suspended in 1 ml PBS and adjusted to 1×10^7^ cells/ml using Vi-Cell Blu cell counter. Aliquots of 1 ml were pelleted and suspended in 1 ml 30 µg/ml Alcian Blue solution and incubated in the dark at room temperature for 15 min. The aliquots were centrifuged and the supernatant was used to quantify Alcian Blue concentration (free/un-bound) at 620nm. This was then used to determine concentration of Alcian Blue dye bound to cells as described previously (Hobson et al. 2004).

### 2.4. Hydrophobicity assay

Cell surface hydrophobicity (CSH) was determined as described by (Rosenberg et al. 1996). Briefly, cells grown overnight were harvested and washed twice with sterile PBS. A cell suspension at OD_600_ of 0.5 was prepared in 3 ml PBS (A0) and was overlaid with 0.4 ml hydrophobic hydrocarbon, n-hexadecane (Sigma-Aldrich). After vigorous vortexing, the two phases were allowed to separate for 10 min at 30°C. The optical density of aqueous phase was measured (A1) and percentage CSH of the eight isolates was determined using following formula: Hydrophobicity (%) = [1-(A1/A0)] × 100.

### 2.5. Adhesion assay

*C. auris* isolates grown overnight were harvested and washed twice with PBS. Cells were then standardized to 2×10^6^ cells/ml in PBS. Adhesion of these isolates was evaluated by using a flow cytometry assay as described by (Silva-Dias et al. 2012). Briefly, 100 µl of the above cell suspension was mixed with 1×10^6^ uncoated carboxylated yellow-green fluorescent polystyrene microspheres (Molecular Probes) and incubated at room temperature for 30 min with rotary shaking at 150 rpm. Following incubation, yeast cell suspensions were vortexed and 30,000 events were analysed using BD Accuri C6 Plus flow cytometer (BD Biosciences, Sydney) – and gated as in (Silva-Dias et al., 2012).

For amyloid exposure, *C. auris* isolates grown overnight were further incubated with 30 µM Thioflavin-T for 1 h and 24 h (Ramsook et al. 2010). Following incubation, cells were harvested, washed and used to quantify mean fluorescence intensity (MFI) using BD Accuri C6 Plus flow cytometer and adhesion assay described respectively. Results for the adhesion assay are expressed as percentage of cells attached to microspheres. Data was collected over three independent experiments with duplicate or triplicate samples, (*n* = 6 or *n* = 9).

### 2.6. Immunofluorescence staining of cell wall with Fc-Pattern Recognition Receptors

The cell wall of eight *C. auris* isolates was incubated with Fc-Pattern Recognition Receptors fusion proteins (Fc-PRRs) to label exposed mannans (CRD4-7-Fc (Mannose Receptor), DC-SIGN and dectin-2), β-glucan (with PRR dectin-1) and exposed chitin (Wheat Germ Agglutinin-FITC) as described by (Vendele et al. 2020). Briefly, 1 × 10^6^ of *C. auris* cells (16 h YPD grown, formaldehyde fixed) were transferred into V-shaped 96-well plates. Plates were centrifuged at 3000 xg for 5 min and supernatants were removed. Cells were incubated with 10 μg/ml dectin-1-Fc in PBS, 1% (v/v) FCS or 10 μg/ml of dectin-2-Fc, CRD4-7-Fc, DC-SIGN-Fc (SinoBiological) in binding buffer (BB) (150 mM NaCl, 10 mM Tris pH 7.4, 10 mM CaCl2 in sterile water + 1% FCS) were then added to appropriate wells and incubated for 45 min on ice. Cells were washed once and then incubated with Alexa Fluor 488 goat anti-human IgG antibody (ThermoFisher) diluted 1/200 in PBS + 1% FCS or BB buffer, or 10 ug/mL Wheat Germ Agglutinin-FITC in HBSS, 1% (v/v) FCS and incubated 30 min on ice. Cells were washed twice before acquired by BD Accuri C6 Plus flow cytometer. Three independent experiments were performed and 30,000 events were recorded for each sample and analysed by FlowJo v10 software.

### 2.7. Scanning electron microscopy

*C. auris* isolates were sub-cultured on YPD agar at 30°C and 37°C for 24 h. Cell colonies intended for ultrastructural analysis were fixed in 2% glutaraldehyde and 2% paraformaldehyde in 0.1 M sodium cacodylate, pH 7.2 for 2 h at room temperature. Colonies were then washed 3 × 5min using cacodylate buffer and subsequently post-fixed with aqueous 1% osmium tetroxide for 1 h. After 3 × 5min washes in deionized water cells were dehydrated in increasing concentrations of ethanol (30%, 50%, 70%, 80%, 90%, 95%, 10 min per step, followed by 2 × 15 min in 100% ethanol), then incubated in 1:1 HMDS (hexamethyldisilazane):ethanol for 1 min and 100% HMDS for 3 min before air drying. Dried colonies were carefully mounted on adhesive carbon tabs applied to aluminium stubs, sputter coated with 10 nm Au/Pd (80/20) and were imaged with a Zeiss GeminiSEM 500 operated between 1.5 – 5 kV using a SE2 detector.

### 2.8. Quantitative PCR

RNA was extracted from *C. auris* isolates, NCPF 8977 (aggregating) and NCPF 13029 (weakly-aggregating) using Monarch RNA extraction kit (NEB cat no. T2010) as per manufacturer’s specification. Prior to extraction, *C. auris* cell pellets were suspended in 800 µl protection buffer (NEB) and 200 µl zirconia beads and lysed by 10 rounds of bead beating at top speed for 30 sec followed by 30 sec of incubation on ice. Samples were then subjected to DNase treatment to remove any residual genomic DNA using Monarch RNA purification kit (NEB cat no. 2040S).

Moloney Murine Leukemia Virus Reverse Transcriptase (M-MLV RT) system for RT-PCR (Promega) was used for cDNA synthesis using 500 ng extracted RNA from all samples. One-tenth volume of cDNA (2 µl) was added to qPCR reactions containing target specific primers (Table 2) and SYBR green master mix (Applied Biosystems) as per manufacturer’s specification. Quantitative PCR was performed on QuantStudio 7 Pro Real Time PCR system (Applied Biosystems). The assay consisted of 10 min denaturation at 95°C followed by 40 cycles at 95°C for 15 sec and 60°C for 1 min. Then, melt curve analysis was performed (60°C to 95°C at a ramp rate of 0.1°C/second). Relative expression of target gene was determined by normalising against housekeeping gene, *ACT1* using following formula:

ΔCt = Ct_target gene_ – Ct_housekeeping gene._ Relative expression = 2_-ΔCt_

### 2.9. DNA extraction and library preparation for Whole Genome Sequencing (WGS)

*C. auris* isolates were sub-cultured in liquid YPD for 24 h at 30°C prior to extraction. DNA was extracted using a phenol-chloroform protocol as described previously (Gautam 2022) and concentration was quantified by Qubit assay kit. Genomic libraries were constructed and sequenced by University of Exeter Sequencing Facility. Tagmentation, PCR amplification and clean-up, library normalisation, pooling and sequencing were carried out on NovaSeq 6000 (Illumina) with 2 × 150 bp paired-end chemistry as previously described (Biswas et al. 2018).

### 2.10. WGS Data Analysis

The Genome Analysis Toolkit (GATK) v.4.1.2.0 (McKenna et al. 2010) was used to call variants. Briefly, raw sequences were pre-processed by mapping reads to the Clade I reference genome *C. auris* 16B25 (Rhodes et al. 2018) using BWA-MEM v.0.7.17 (Li 2013). Next, duplicates were marked, and the resulting file was sorted by coordinate order. Intervals were created using a custom bash script allowing parallel analysis of large batches of genomics data. HaplotypeCaller was executed in GVCF mode with the haploid ploidy flag. Variants were imported to GATK 4 GenomicsDB and hard filtered if QualByDepth (QD) < 2.0, FisherStrand (FS) > 60.0, root mean square mapping quality (MQ) < 40.0, Genotype Quality (GQ) ≥ 50, Allele Depth (AD) ≥ 0.8, or Coverage (DP) ≥ 10.

To identify aneuploid chromosomes, depth of coverage was calculated for each of the samples. Sorted BAM files prepared in the pre-processing phase of SNP calling were passed to Samtools v.1.2 (Li et al. 2009) and pileup files were generated. Read depth was normalised by total alignment depth and plotted against the location in the genome using 10 kb non-overlapping sliding windows.

### 2.11. Phylogenetic and evolutionary analysis

To construct species-specific phylogenetic trees, all sites that were either a homozygous reference or SNP in every isolate were identified using ECATools (https://github.com/rhysf/ECATools) and concatenated into a FASTA file. Our unrooted tree included 202,878 phylogenetically informative sites. Phylogenetic trees were constructed with RAxML PThreads v.7.7.8 (Stamatakis 2006) using the general-time-reversible model and CAT rate approximation and midpoint rooted.

All *C. auris* protein sequences were screened for GPI anchors using PredGPI, (Pierleoni, Martelli, and Casadio 2008) identifying 68 genes with high probability of encoding a GPI anchor, 27 with a probability, 20 with a weak probability, and the remaining protein-encoding genes of *C. auris* (*n* = 5,153) not predicted to encode a GPI anchor. Enrichment of P/A polymorphisms (*n* = 71) that had high probability of encoding a GPI anchors (*n* = 11) was assessed using the R phyper statistical test where q = 11, m = 68, n = 5200, k = 71, and lower.tail = FALSE. Secretion signals were predicted using SignalP4 (Petersen et al. 2011).

The direction and magnitude of natural selection for each isolate were assessed by measuring the rates of non-synonymous substitution (*dN*), synonymous substitution (*dS*) and omega (ω = *dN*/*dS*) using the yn00 program of PAML (Yang 2007), which implements the Yang and Nielsen method, taking into account codon bias (Yang and Nielsen 2000). The program was run on every gene in each isolate using the standard nuclear code translation table.

## 3 Results

### 3.1. Aggregation in C. auris clinical isolates

We first confirmed and assessed the extent of aggregation variability in eight clinical isolates of *C. auris* (Table 1), which is a property that is suspected to contribute to virulence and immune evasion strategies (Borman, Szekely, and Johnson 2016). Four of the eight isolates exhibited a strong aggregating phenotype (Fig. 1) while others were weakly-aggregating in liquid YPD. As previously reported by Borman *et al*., the aggregates of the four strong aggregating isolates were resilient to physical disruption by vortexing.

**Fig. 1.**
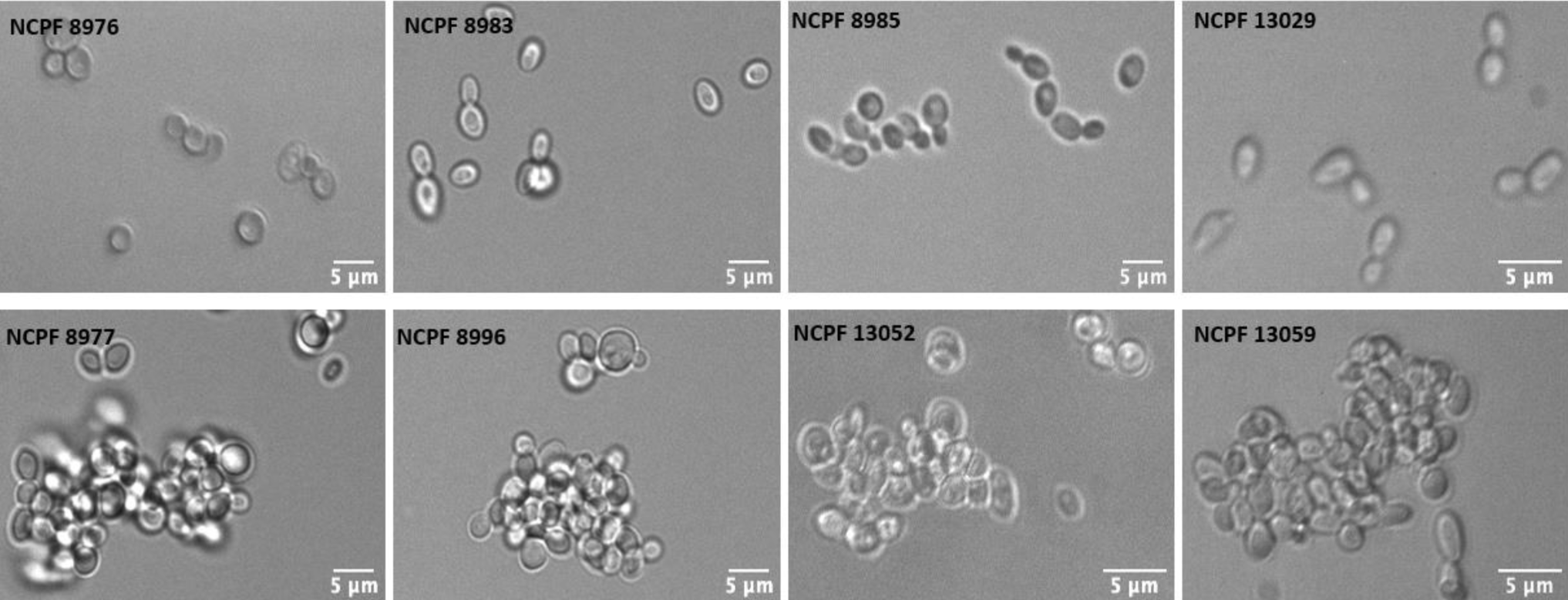
DIC images of weakly-aggregating (upper panel) and strongly aggregating isolates (lower panel) of *C. auris* in PBS suspension. All isolates were grown on YPD agar for 24 h at 30°C before imaging.

### 3.2. Genomic analysis of the eight C. auris clinical isolates to understand aggregation

Phylogenetic analysis determined that the four aggregating strains belonged to Clades III and IV, while the four weakly-aggregating strains are from Clades I and II. Given that aggregation is not clade-specific (Chakrabarti and Sood 2021), we hypothesised that the aggregating phenotype may relate to multiple independently evolving gene sets, allelic and possible epigenetic traits.

To explore the genetic differences between strongly-aggregating and weakly-aggregating isolates, we undertook an *in silico* screen of polymorphic genomic regions with an emphasis on genes encoding secreted proteins. Depth of coverage plots indicated that there were few obvious chromosomal copy number variation (CNV) (Fig. 2C). The few examples of CNV included scaffold 8.16 in isolate NCPF 8996, as well as non-uniform coverage in several isolates (NCPF 13029, NCPF 13059) likely owing to common technical library preparation issues biasing for increased coverage at telomeric regions.

**Fig. 2.**
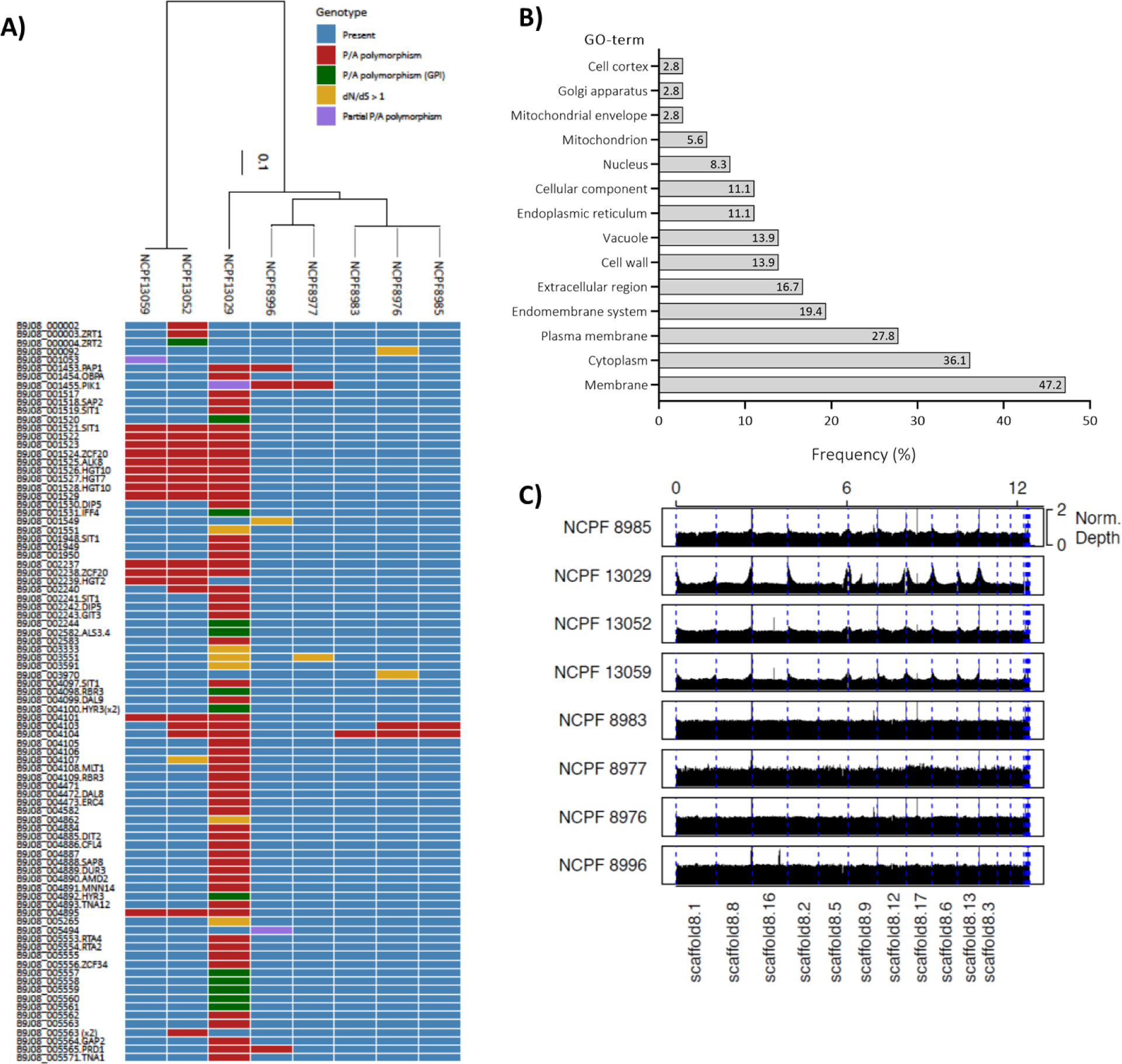
Genome dynamics and repertoire of genes in *C. auris* isolates. The weakly-aggregation strains are: NCPF 8976, NCPF 8983, NCPF 8985 and NCPF 13029 and the strongly-aggregating strains are: NCPF 8977, NCPF 8996, NCPF 13052 and NCPF 13059. (A) Strain-specific differences in presence (blue) or absence (red) of genes is highlighted. Gene deletions that are predicted to be GPI-anchor protein encoding genes in the Clade I reference strain (green), genes with signatures of positive selection (d_N_/d_S_ > 1; yellow) and partial P/A polymorphisms purple) are also highlighted. The Locus ID is provided for all genes with these differences across the 8 isolates. Common gene names are provided where known. A phylogenetic tree demonstrating the placement of each isolate into the 4 clades was constructed with RAxML using the GTR model and CAT rate approximation and shown at the top of the heatmap. (B) GO term analysis of all genes absent in weakly-aggregating isolates. (C) Chromosomal position and gene number across chromosomes in the genomes for the eight clinical isolates. Depth of coverage plots show no chromosomal copy number variation.

We identified 79 gene presence/absence polymorphisms (P/A polymorphisms; defined as having breadth of coverage < 10% across the gene length) in one or more strains (Fig. 2A, B). None of the P/A polymorphisms were found exclusively in strongly aggregating isolates or weakly aggregating isolates, and none of the isolate’s genomes lacked essential genes encoded by the Clade I reference genome involved in budding, cell division or separation, suggesting that gene loss is not the cause of aggregation across these strains.

Many of the P/A polymorphisms were due to deletions identified in several isolates. For example, isolates NCPF 13052 and NCPF 13059 had a 20 kb deletion from the end of scaffold 8.5 (corresponding to chromosome 3 in the B8441 genome assembly, from approximately positions 1,021,142 - 1,041), encompassing 9 genes. Isolate NCPF 13029 had the same deletion, although it was 32 kb (beginning at approximately position 1,009,684) and encompassing 15 genes including 2 for proteins of predicted GPI anchors (IFF4 and B9J08_001520). In total, all P/A polymorphisms identified across the 8 isolates could be grouped into approximately 27 larger deletions. Other notable regions that were absent in >1 isolates included the start of chromosome 6 till position 21,267, the start of chromosome 8 till position 33,225, the end of chromosome 9 (starting at position 960,751) and chromosome 12 (positions 26,146 till 46,320). Together, these 5 genomic regions on chromosomes 3, 6, 8, 9 and 12 were responsible for 63% (*n* = 50/79) P/A polymorphisms identified.

Several P/A polymorphisms had an association with adhesion ability in individual isolates. For example, most (*n* = 71/79; 90%) of the P/A polymorphisms were identified in the weakly-aggregating Clade II isolate NCPF 13029, 15% (*n* = 11/71) of which were for genes that had a high probability of encoding a predicated GPI anchor, including *ALS4* which is known to be involved in adhesion (Hoyer et al. 2008; Zhao et al. 2005) (Fig. 2A). Noteworthy, this weakly-aggregating isolate did not even form small aggregates of <4 cells. Given that only 68 of the 5,268 predicted protein encoding genes in *C. auris* had a high probability of encoding a GPI anchor, this represents a significant enrichment among those gene deletions (Hypergeometric test *p*-value = 5.51E^-11^). GO-term analysis of these P/A polymorphisms revealed that 77.8% were associated with cell wall, membrane and extracellular proteins. Together, this data suggests that loss of aggregation in isolate NCPF 13029 is likely caused by the loss of one or more cell surface adhesins rather than a defect in cell separation.

To explore if signatures of positive selection underlie the aggregation phenotype, we identified genes with signatures of positive selection (*dN*/*dS* > 1) compared with the Clade I reference strain, identifying only 10 genes in total (Fig. 2A). Again, most of these genes (*n* = 6/10) were in isolate NCPF 13029 and were mostly poorly characterised (all lacked GO-terms, and none have previously been reported to be involved in aggregation). Therefore, this approach has not identified any gene candidates for aggregation, although those genes may be interesting candidates for future experiments. We have not ruled out the possibility of key mutations driving aggregation phenotypes.

### 3.3. Expression of chitinases and chitin synthases in strongly aggregating and weakly-aggregating isolates

Since dysregulation of the chitinase gene *CTS1* is suspected to cause aggregation (Santana and O’Meara 2021), we compared expression of chitinase genes *CTS1* and *CTS2* in the strongly-aggregating isolate NCPF 8977 with the weakly-aggregating isolate NCPF 13029. We also compared expression of chitin synthases (*CHS1, CHS2, CHS3, CHS4* and *CHS8*) that regulate septum formation leading to cell separation. We found that expression levels of *CTS1* were comparable between the strongly aggregating and weakly-aggregating isolate. However, expression of *CTS2* was substantially lower in the strongly aggregating isolate compared to the weakly-aggregating isolate (Fig. 3A). Additionally, the strongly aggregating isolate expressed substantially lower expression of *CHS1* and *CHS2* compared to the weakly-aggregating isolate. The *CHS3* mRNA level was low in the weakly-aggregating isolate but higher in the aggregating isolate (Fig. 3B). Therefore, the transcript expression of most genes involved in the formation and dissolution of the chitinous septum formation was lower in the strongly-aggregating isolate than the weakly-aggregating isolate.

**Fig. 3.**
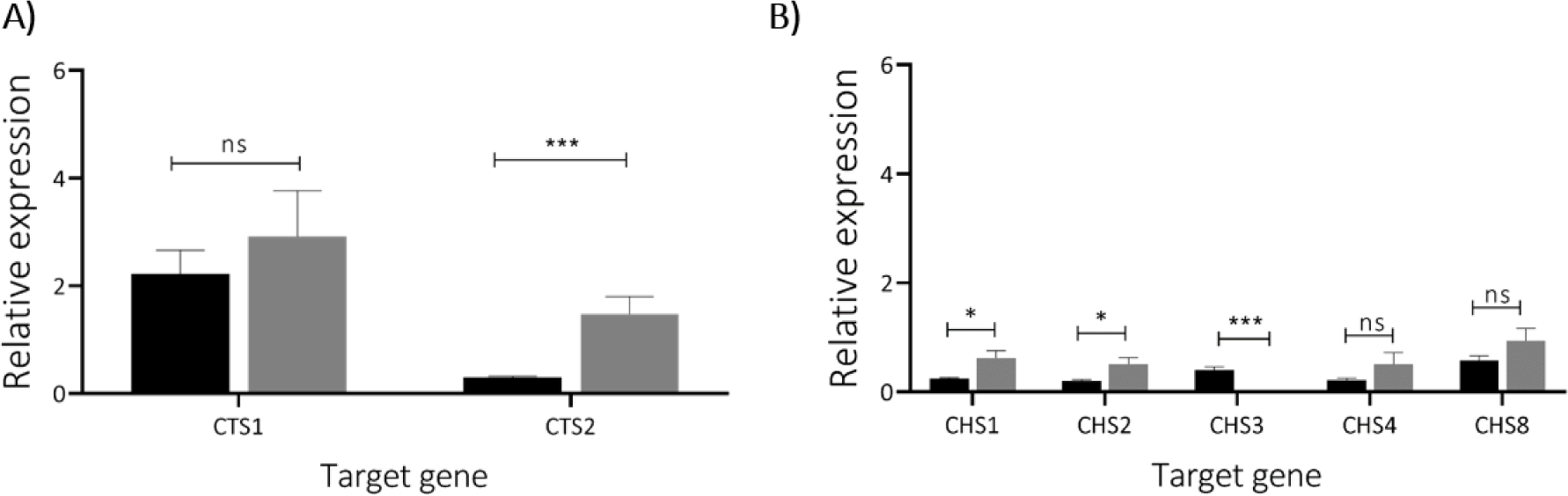
Expression of (A) chitinases and (B) chitin synthases in strongly aggregating isolate NCPF 8977 (black bars) and weakly-aggregating isolate NCPF 13029 (grey bars). Relative expression was determined by normalization against housekeeping gene *ACT1*, levels of which are comparable in both strains. Data obtained from three independent experiments performed in triplicates. Error bars represent standard deviation. Student t-test used for statistical analysis. *** represent *p*-values < 0.001 (where *n* = 9).

### 3.4. Cell separation in C. auris clinical isolates

We investigated if temporal differences in cell separation or defects in separation of daughter cells from mother cells explained the aggregation phenotype by using a combination of microscopy and microfluidics. Individual yeast cells from strongly aggregating and weakly-aggregating isolates were trapped in CellASIC microfluidic system and growth was assessed for 8 h. Each isolate was grown in the presence of enriched YPD medium supplemented with 1 µg/ml Calcofluor White (CFW) to visualise cell wall chitin and the chitin rich septal wall separating a mother and daughter cell. The architecture of septal rings and unipolar cell growth of daughter cells emerging from mother cells was followed. Over a period of 8 h, normal chitin-rich septal rings were observed in all dividing *C. auris* isolates including aggregating strains (Fig. 4). The final step in yeast cell separation is marked by degradation of chitinous primary septum. For all tested isolates, a reduction in Calcofluor White fluorescence was observed at all bud sites as a daughter cell separated from the mother cell suggesting cell separation by chitin dissolution still occurred. Following cell separation, daughter cells remained in close vicinity to mother cells (Supplementary movie S1). Contrary to previous hypothesis (Santana and O’Meara 2021) and despite minor changes in the expression of genes forming and degrading chitin, an obvious cell separation defect was not observed in any of the strongly aggregating or weakly-aggregating isolates.

**Fig. 4.**
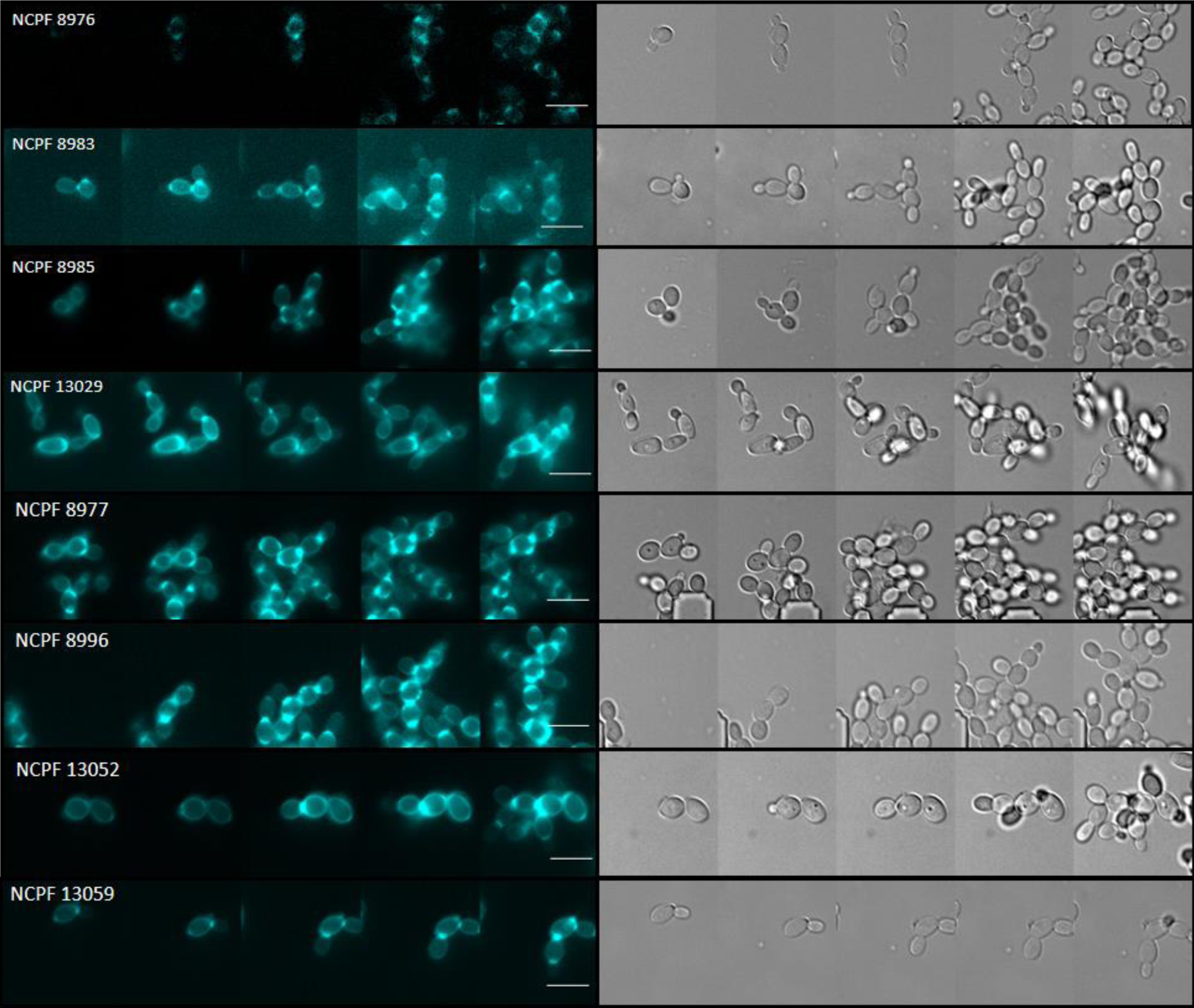
Cell separation in *C. auris* isolates in microfluidic growth chambers. Time lapse montage of individual isolates stained with 1 µg/ml Calcofluor White (left panel) to label cell wall chitin, complimented with DIC images (right panel). Calcofluor White staining cell wall and chitin rich septa seen in all, strongly-aggregating and weakly-aggregating isolates. Images taken using DeltaVision widefield fluorescence microscope. Scale bar represents 5 µm.

### 3.5. Adhesion and aggregation profile of strongly aggregating and weakly-aggregating C. auris isolates

Cell surface adhesins are primarily responsible for adhesion in *Candida* species. Increased adhesion capacity in some *C. auris* isolates could be responsible for the aggregation phenotype (Klotz et al. 2007; Bing et al. 2023). Hence, the adhesion of strongly-aggregating and weakly-aggregating isolates to polystyrene microspheres was determined. We found that adhesion capacity of strongly aggregating isolates was comparable to the weakly-aggregating isolates (Fig. 5A). The NCPF 8996 aggregate forming isolate had the lowest adhesion capacity (65.5% of cells attached to microspheres) of the eight tested isolates. Therefore, there was no correlation between adhesion capacity and aggregation since high and low adhesion capacity was found in both strongly aggregating and weakly-aggregating strains. However, the adhesion capacity of *C. auris* isolates was substantially greater (ranging from 65.5% to 94.4% cells attached to microspheres) than the *C. albicans* reference strain SC5314 (25.4% attached) and other *Candida* species tested previously (Silva-Dias et al. 2015).

**Fig. 5.**
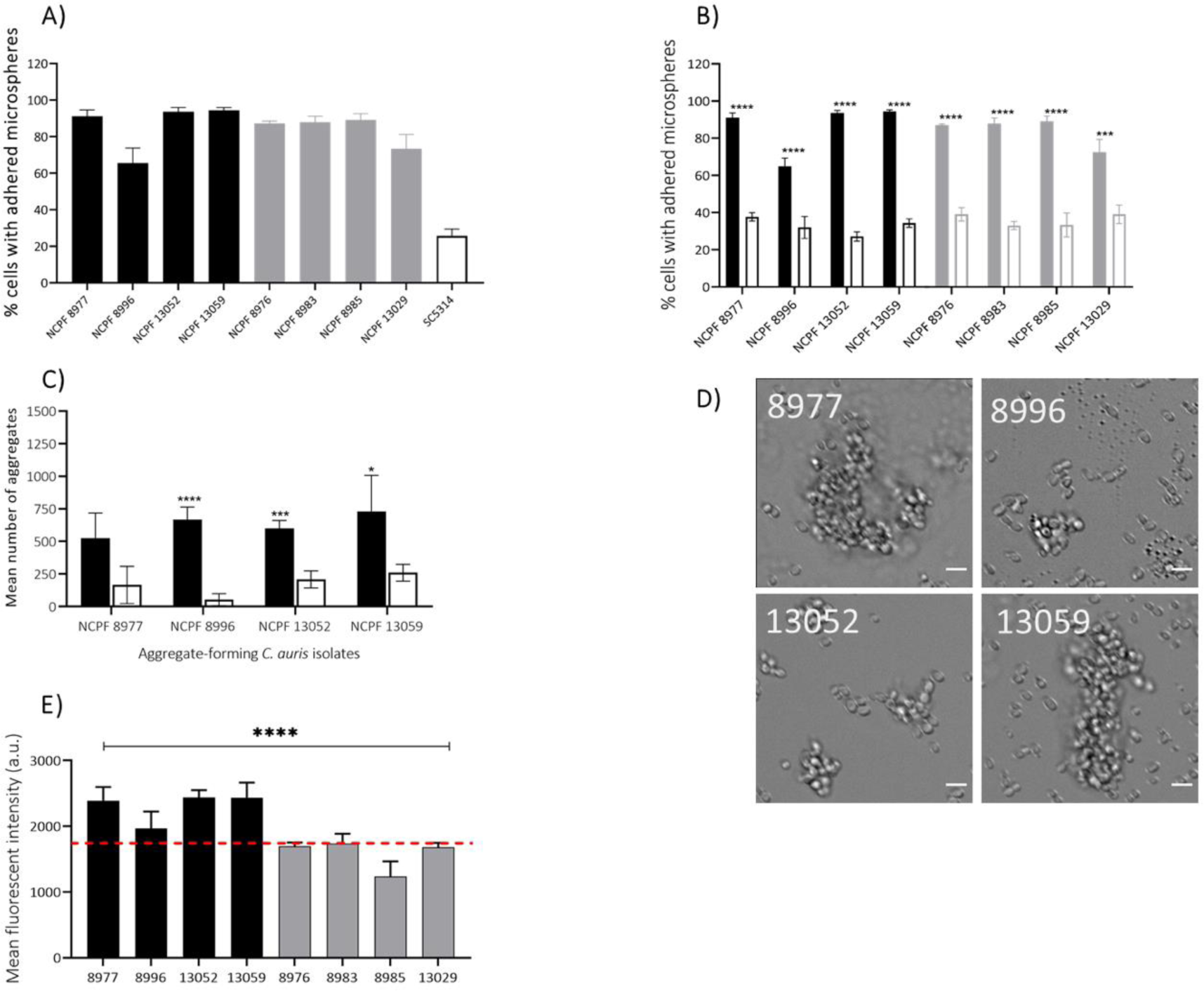
The lack of correlation between adhesion and aggregation of *C. auris* strains. (A) Percentage of cells with adhered microspheres. Adhesion capacity of strongly-aggregating *C. auris* isolates (black) compared with weakly-aggregating isolates (grey) and *C. albicans* SC5314 (white bar, black-border) following incubation with uncoated carboxylated yellow-green fluorescent polystyrene microspheres for 30 min. (B) Percentage of cells adhered to microspheres following treatment with 30 µM Thioflavin-T for 24 h and used to determine adhesion capacity as before (solid bars = untreated, white bars with black border = treated cells). (C) Mean number of aggregates in strongly-aggregating isolates following 24 h incubation with Thioflavin-T. Data obtained from 60 images per isolate over 3 independent experiments. (D) Representative images of aggregating strains after 24 h incubation with Thioflavin-T. Images acquired using DeltaVision widefield fluorescence microscope at 200× total magnification. Scale bar represents 5 µm. (E) Amyloid staining (FITC channel, BD Accuri) following incubation with 30 µM Thioflavin-T for 1 h at 30°C. Red horizontal line represents highest MFI recorded among weakly-aggregating isolates. Graphs obtained from 3 independent experiments performed in duplicates (E); (n=6) and triplicates (n=9). Error bars represent standard deviation. Student-T test used for statistical analysis. P****<0.00001, P**<0.01, P*<0.05, (n=9).

### 3.6. Inhibition of amyloid protein function attenuates adhesion and aggregation

Many adhesins possess amyloid-forming sequences that can be hypothesised to make a contribution the capacity for aggregation. Thioflavin-T binds and inhibits surface amyloids and therefore may also inhibit aggregation. Therefore, *C. auris* isolates were incubated with 30 µM Thioflavin-T for 1 h and 24 h. We quantified surface amyloids in all isolates, and assessed adhesion following inhibition of surface amyloids and quantified microscopically the aggregation of isolates exposed to Thioflavin-T. We discovered that all aggregating isolates had greater staining for surface amyloid proteins compared to weakly-aggregating isolates (Fig. 5E). Following incubation with Thioflavin-T for 24 h, the capacity of these cells to adhere to polystyrene microsphere was also reduced (Fig. 5B). As expected, adhesion of Thioflavin-T treated isolates to polystyrene microspheres was substantially lower than their untreated counterparts. An average 2 fold decrease was observed for all tested *C. auris* isolates following Thioflavin-T treatment (average % adherent cells = 85% before treatment and 34.4% after treatment). Furthermore, the aggregating isolate NCPF 13052 showed a 3.47 fold decrease (from 93.7% to 27.2%) in adhesion following Thioflavin-T treatment. However, microscopy revealed that even after 24 h of incubation with Thioflavin-T, aggregating-isolates continued to form aggregates (Fig. 5C, D). Aggregation was attenuated for strongly-aggregating isolates under the conditions tested, but was not completely suppressed. These observations suggest that amyloid formation is important for aggregation but that other unidentified factors may also contribute to aggregation.

### 3.7. Cell surface charge and hydrophobicity in C. auris isolates

We investigated the effects of cell surface hydrophobicity and charge on aggregation. Cell surface hydrophobicity has previously been shown to improve adhesion and aggregation in yeasts (Silva-Dias et al. 2015; Yu-Lai Jin 2001). To determine if surface hydrophobicity contributes to aggregation in *C. auris*, the eight isolates were incubated in the hydrophobic solvent hexadecane for 10 min. However, no link was observed between aggregation and their relative hydrophobicity (Fig. 6A). Two of the four strongly aggregating isolates, NCPF 13052 and NCPF 13059 had the highest percent of cell surface hydrophobicity (34.2% and 34.6%, respectively). However, a strongly aggregating (NCPF 8996) and weakly-aggregating (NCPF 13029) isolate had the lowest percent of cell surface hydrophobicity (20.7% and 10.4%, respectively). All other isolates (the strongly-aggregating-NCPF 8977; weakly-aggregating NCPF 8983 and 8985), exhibited comparable cell surface hydrophobicity.

**Fig. 6.**
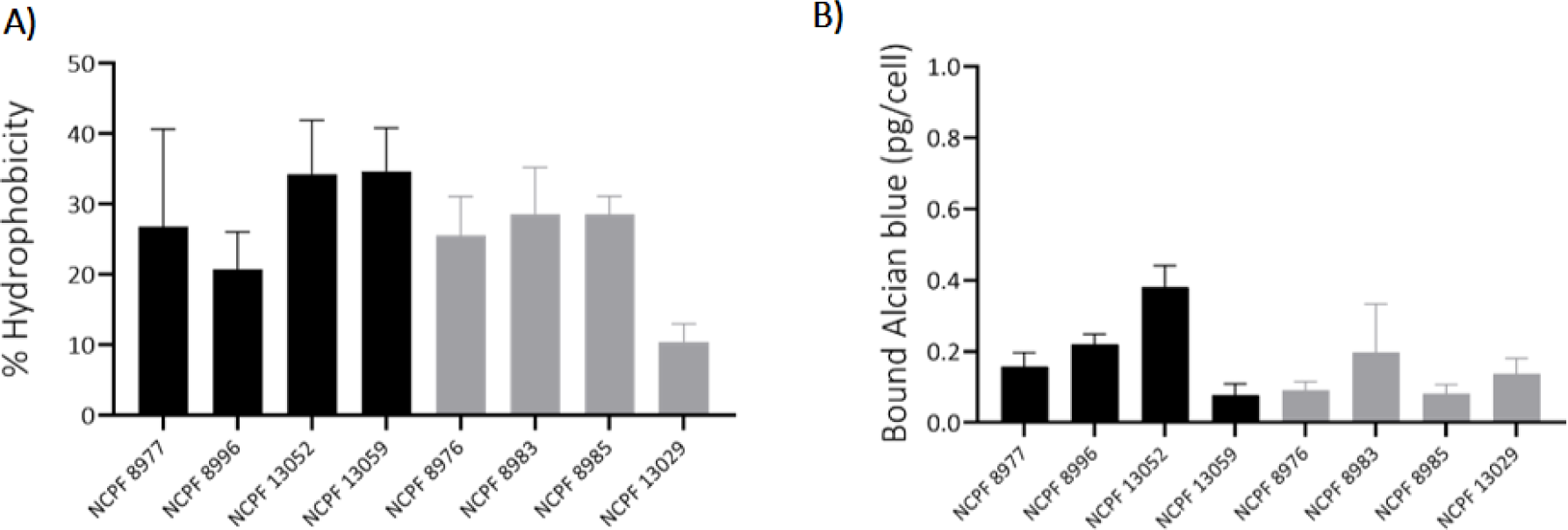
Cell surface properties (hydrophobicity and cell charge) of *C. auris* isolates. (A) Cell surface hydrophobicity assessed using microbial adhesion after hydrocarbon treatment. (B) Cell surface charge assessed using 30 µg/ml Alcian Blue cationic dye. Graphs represent means of aggregating (black bars) and weakly-aggregating (grey bars) isolates. Experiments based on 3 independent experiments in triplicate; error bars represent standard deviation (*n* = 9).

Net cell negative surface charge can contribute to cell aggregation (Wilcocks and Smart 1995). Hence, surface charge of the eight *C. auris* isolates was determined by staining with 30 µg/ml of the cationic Alcian Blue dye that in *Candida* cells is known to bind predominantly to phosphomannan in the outer cell wall layer (Hobson et al. 2004). Higher Alcian Blue binding would be indicative of a higher net negative charge on cell surface. As seen with hydrophobicity, no link was found between the aggregation phenotype and surface charge (Fig. 6B). Strongly-aggregating isolates, NCPF 13052 and NCPF 13059 exhibited the highest Alcian Blue binding (0.38 pg/cell) and lowest binding (0.08 pg/cell) respectively.

### 3.8. Cell wall composition and carbohydrate content in C. auris isolates

We examined the global cell wall composition of the eight *C. auris* isolates by measuring their access to a range of carbohydrate recognising pattern recognition immune receptors for β-glucan (dectin-1-Fc), exposed chitin (WGA-FITC) and mannans (dectin-2-Fc, CRD4-7-Fc and DC-SIGN-Fc). Mannan-recognising PRR-Fc’s demonstrated differential binding across all eight isolates (Fig. 7A). For example, the weakly-aggregating isolate NCPF 8976 demonstrated strong dectin-2-Fc binding. However, dectin-2-Fc binding in weakly-aggregating isolate NCPF 8983 was comparable to strongly-aggregating isolates NCPF 13052 and NCPF 13059. Similarly, all *C. auris* isolates showed a range of binding profiles to DC-SIGN-Fc and CRD4-7-Fc with no overall correlation to aggregation. There was no correlation with the degree of exposed chitin content (WGA-FITC binding) in the cell walls of the isolates. Although dectin-1-Fc binding was generally stronger in strongly-aggregating isolates, its binding to the weakly-aggregating isolate NCPF 13029 was comparable to strongly-aggregating strains.

**Fig. 7.**
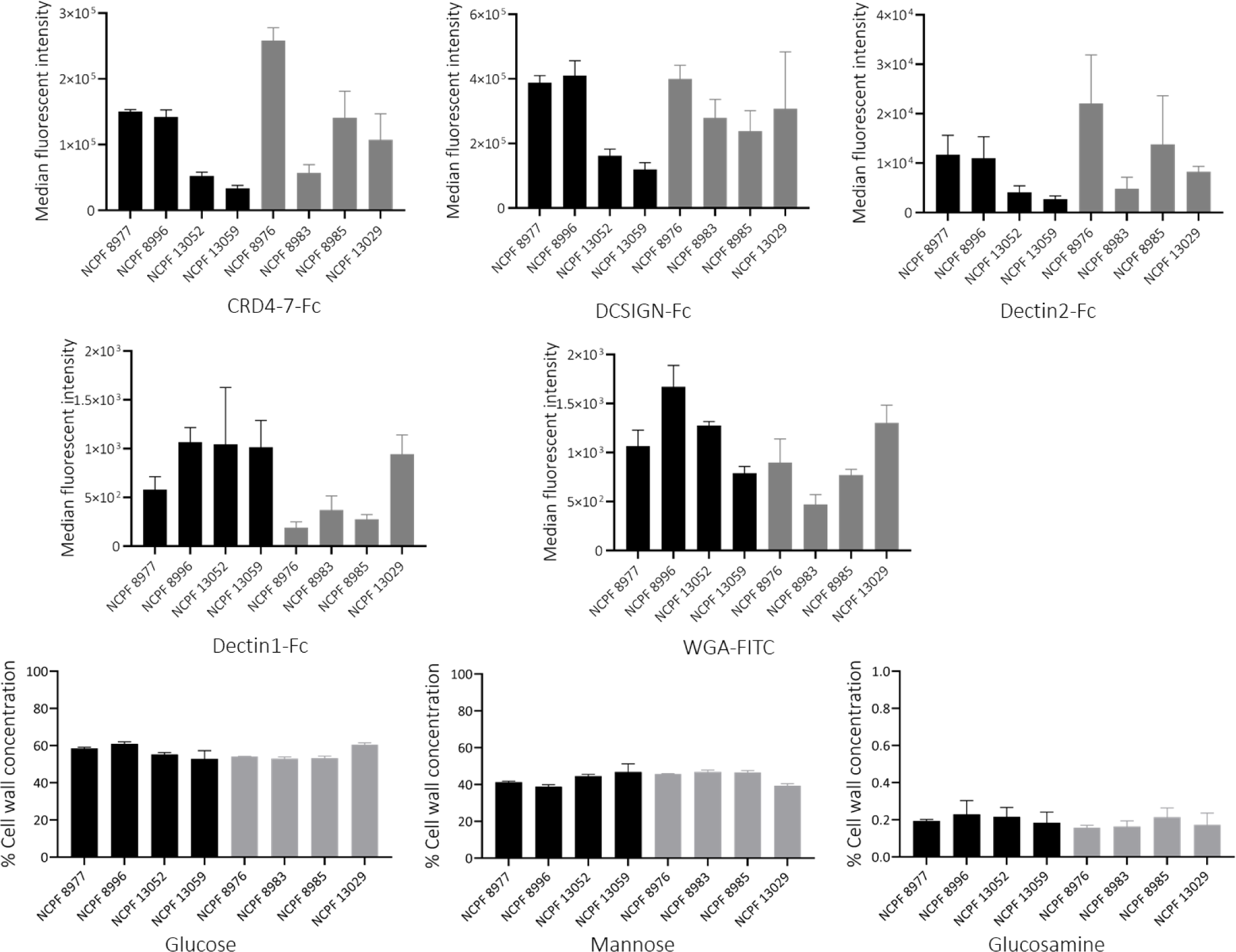
Cell wall properties based on binding to Pattern Recognition Receptors (PRRs) and carbohydrate analyses of the eight *C. auris* isolates. (A) Binding of Fc-PRRs to *C. auris* isolates. Fc-PRR binding in strongly aggregating (black bars) and weakly-aggregating (grey bars) isolates presented as median fluorescence intensities (MFI) of probe binding. All isolates were grown in YPD for 16 h at 30°C before fixation and staining. Experiments were performed in triplicate, error bars represent standard deviation. (B) Relative cell wall polysaccharide composition in *C. auris* isolates. Cells were grown overnight in YPD at 30°C and cell wall was extracted and hydrolysed by trifluoroacetic acid and the hydrolysate analysed for monosaccharides by HPIC. Black bars represent aggregating isolates and grey bars represent weakly-aggregating isolates. Experiments were performed in triplicates, error bars represent standard error (*n* = 9).

The percent of principle carbohydrates (glucose, glucosamine and mannose) that make up the cell wall polysaccharides was determined using High Pressure Ion Chromatography (Fig. 7B). We discovered that despite each isolate having differential Fc-PRR probe binding, the percent of glucose, mannose and glucosamine in the cell wall of all eight isolates were broadly comparable to each other and no statistically significant differences were observed for any of these basic carbohydrates between strongly aggregating and weakly-aggregating isolates. This experiment suggested that although isolate specific differences were observed in the level and organization of cell wall exposed β-glucan, exposed chitin and mannans, these differences are unlikely to explain the aggregation phenotype.

### 3.9. Effects of temperature on aggregation

Three of the four weakly-aggregating isolates demonstrated some increased tendency for aggregation when incubated at 37°C. This led us to investigate the effects of temperature on aggregate formation in *C. auris* isolates. The eight isolates were pre-grown at 30°C and further incubated at 30°C and 37°C for 24 h. All tested isolates displayed a substantial increase in the number of aggregates formed at 37°C compared to 30°C (Fig. 8). NCPF 13029 and NCPF 8985, were very weakly aggregating at 30°C also exhibited some degree of aggregation at 37°C.

**Fig. 8.**
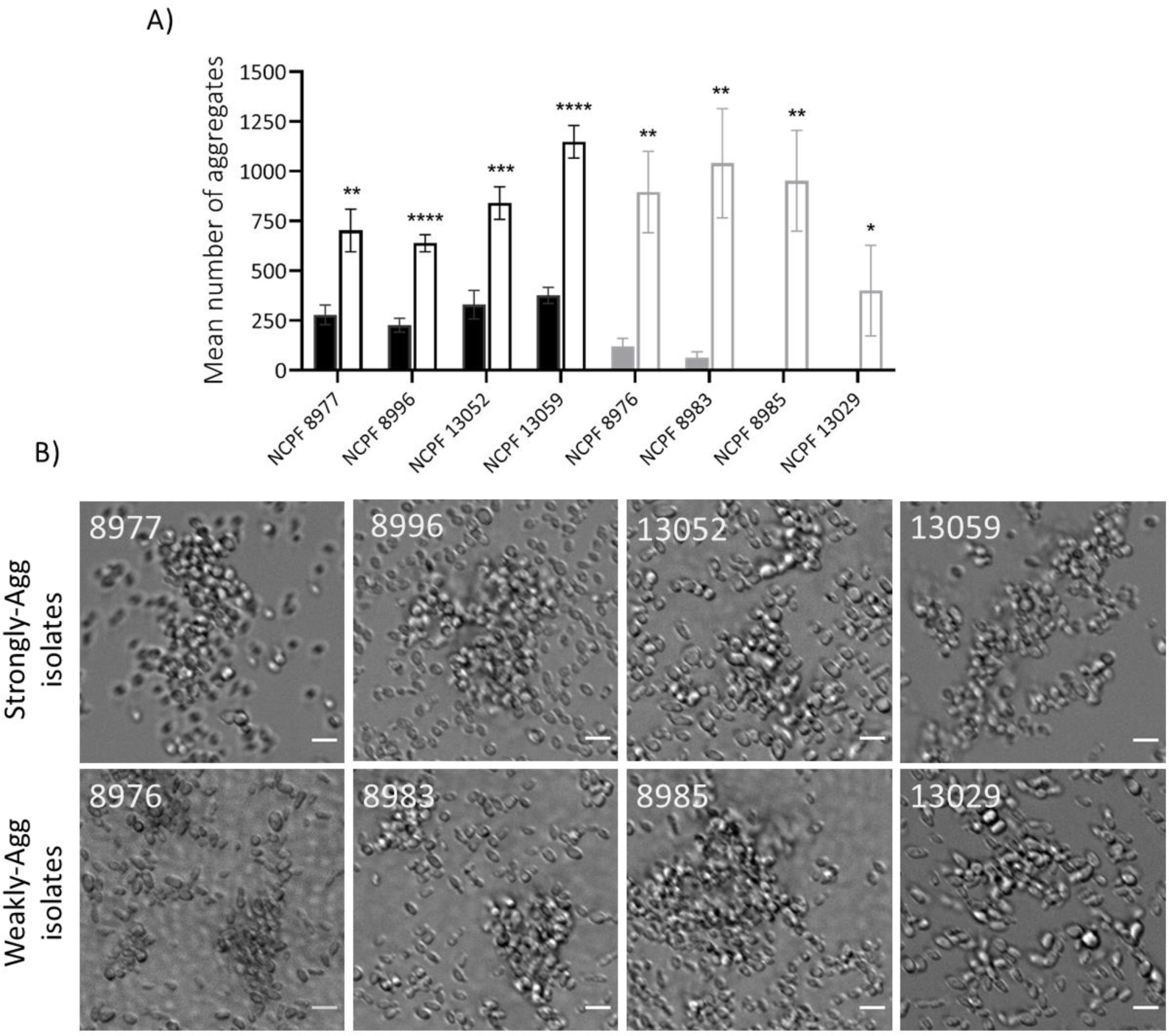
Effects of temperature on aggregation in *C. auris*. (A) Quantification of average number of aggregates in *C. auris* isolates following 24 h incubation at 37°C on YPD agar. Solid bars represent samples incubated at 30°C and lined bars represent samples incubated at 37°C. (B) Representative images of *C. auris* isolates following 24 h incubation at 37°C on YPD agar. Images acquired using DeltaVision widefield fluorescence, scale bar represents 5 µm. Data obtained from 60 images per isolate over 3 independent experiments. Error bars represent standard deviation. Student t-test used for statistical analysis P****<0.00001, P***<0.001, P**<0.01, P*<0.05, (*n* = 9).

### 3.10. Expression of genes involved in biofilm formation

Given the enhanced aggregation capacity of all isolates at 37°C compared with 30°C, we hypothesised that 37°C might trigger genes involved in morphogenesis or biofilm formation that in turn might influence aggregation. Previous studies have shown that genes involved in morphological transition and ECM formation (e.g., *BIG1, EFG1, TEC1* and *XOG1*) confer hyper-adherent properties to *C. albicans* cells (Umeyama et al. 2006; Li and Palecek 2003; Sonneborn, Tebarth, and Ernst 1999; Lohse and Johnson 2010; Schweizer et al. 2000; Nobile and Mitchell 2005; Taff et al. 2012). We hypothesized that increased expression of such protein(s) could contribute to aggregation in *C. auris*. Therefore, we determined expression of these crucial genes involved in morphological changes and biofilm formation. Of the tested target genes, expression of *XOG1* was greater in strongly-aggregating isolate (NCPF 8977; 1.47x compared to weakly-aggregating isolate (NCPF 13029; 0.66) (Fig. 9). Other tested target genes did not show a substantial difference in expression between the two isolates. Furthermore, expression of *XOG1* was substantially greater in the strongly-aggregating isolate NCPF 8977 at 37°C (4.12X) compared to that at 30°C (1.47X). It is therefore possible that expression differences of Xog1 β-exoglucanase contributes to aggregation.

**Fig. 9.**
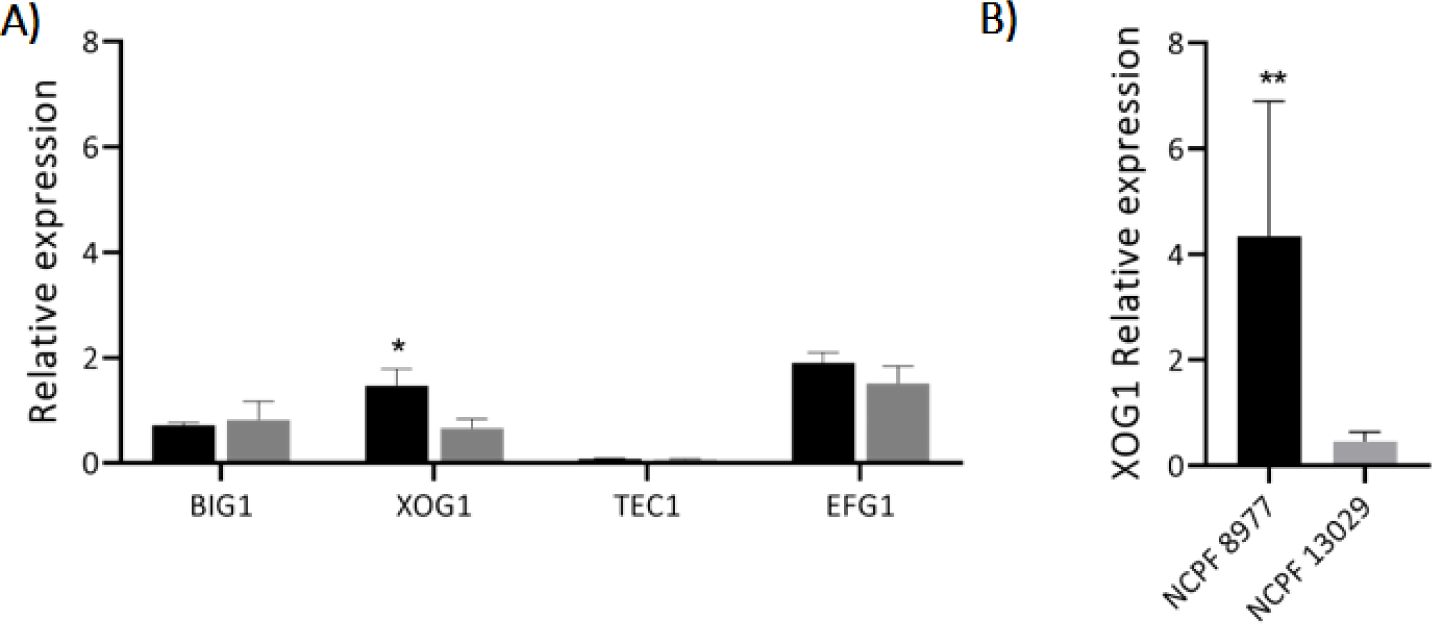
Expression of (A) morphogenesis associated genes in strongly aggregating isolate NCPF 8977 (black bars) and weakly-aggregating isolate NCPF 13029 (grey bars). (B) Expression of *XOG1*, exoglucanase enzyme implicated in ECM formation in biofilms in strongly-aggregating and weakly-aggregating isolates grown at 37°C. Relative expression was determined by normalization against housekeeping gene *ACT1*, levels of which were similar in both strains at 30°C and at 37°C. Data obtained from 3 independent experiments performed in triplicates. Error bars represent standard deviation. Student t-test used for statistical analysis. P**<0.01, P*<0.05 (*n* = 9).

### 3.11. Extracellular material production in aggregating isolates

We examined *extracellular material (*ECM) formation in the eight strains grown at 30°C and 37°C. Small amounts of ECM were seen between adjacent cells of aggregating isolates that had been grown at 30°C which displayed a rough cell surface (Fig. 10). The amount of ECM observed on SEMs varied between aggregating isolates. The aggregating strains NCPF 13052 and NCFP 13059 appeared to be coated in a ‘blanket-like’ layer of ECM. Conversely, aggregating strains NCPF 8977 and NCPF 8996 displayed ECM that was unevenly distributed on the cell surface and was more concentrated where yeast cells adhered to each other. At 37°C, ECM was imaged in all eight isolates to varying degrees thus corroborating previous observations of increased aggregation of “weakly-aggregating” isolates at higher temperature. Based on these findings the production of ECM may also contribute to aggregation.

**Fig. 10.**
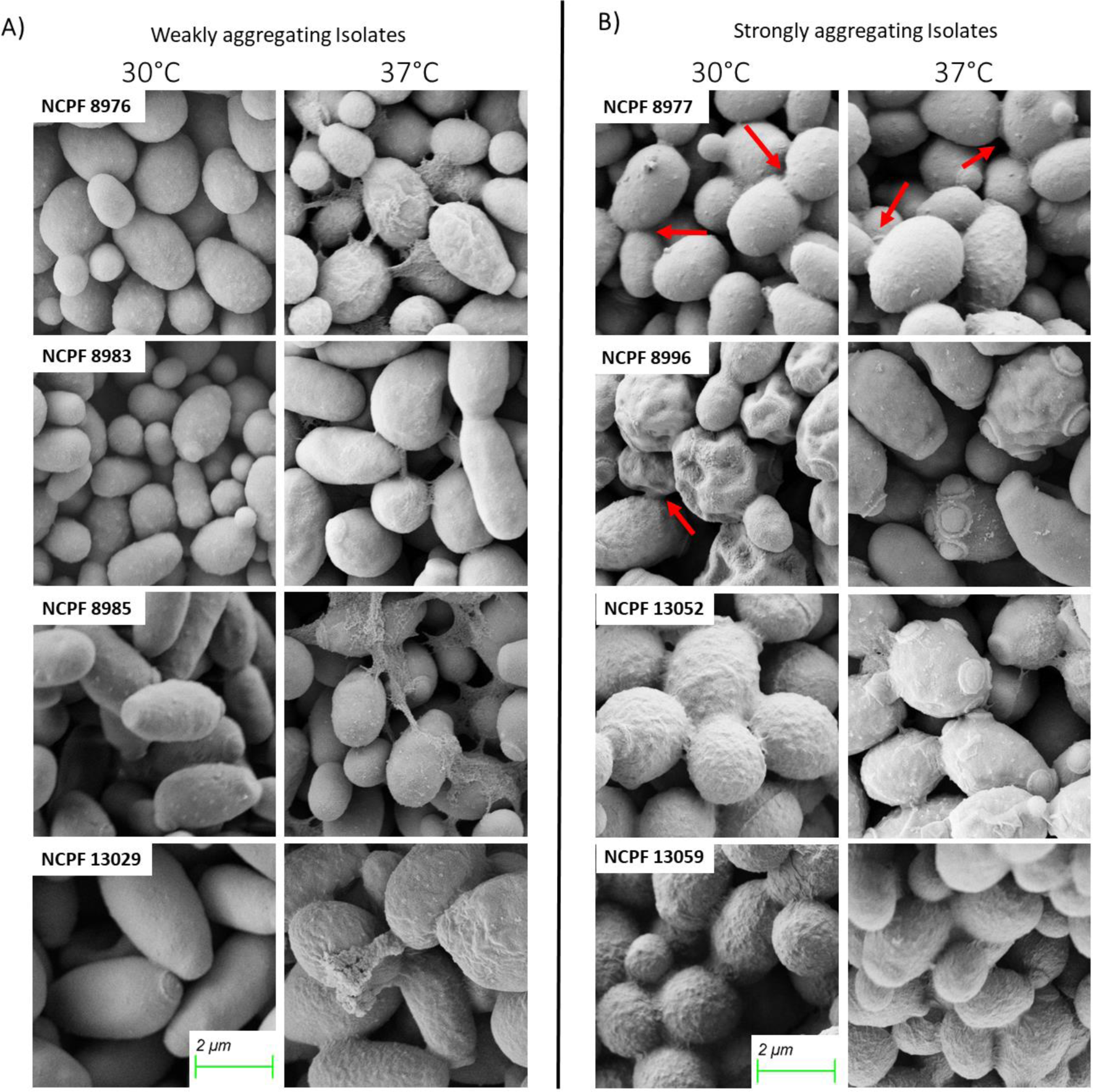
Scanning electron microscopy (SEM) of *C. auris* showing ECM associated with aggregated cells. (A) weakly-aggregating isolates and (B) strongly-aggregating isolates grown at 30°C and 37°C. The cells had varying amounts of ECM which was more prominent at 37°C than at 30°C. Red arrows indicate areas of surface or extracellular deposits suspected to play a role in aggregation. Scale bar, 2 µm.

## 4 Discussion

Strains of many *Candida* species are known to form aggregates; usually under certain environmental or stressful conditions. For example, hypha formation of *C. albicans* is associated with very strong aggregation of germ tubes (Kumar et al. 2015; Mukaremera et al. 2017) and caspofungin treatment can induce cell-aggregation (Gregori et al. 2011). Beads coated with bovine serum albumin or fibronectin can cause *C. albicans* to form homotypic aggregates with other *C. albicans* cells and heterotypic aggregates with *C. glabrata* and *C. parapsilosis* (Klotz et al. 2007). Furthermore, growth in the presence of acetic acid has also been shown to induce aggregation in *C. glabrata* yeast cells in a biofilm (Mota et al. 2015). However, unlike environment-induced aggregation, recent reports have described strain-specific aggregating phenotypes in *C. auris* that are linked with colonisation *in-vivo* and lower pathogenicity in a systemic model of infection compared to weakly-aggregating *C. auris* isolates (Borman, Szekely, and Johnson 2016). Because of its clinical importance, understanding the mechanism(s) of aggregation in *C. auris* has attracted widespread interest in recent years. As a result, two hypotheses have emerged which are investigated in this study – namely that 1) differences in aggregation are due to defects in cell separation and 2) cell surface properties and protein(s) mediate cell-to-cell adhesion.

Microscopy of *C. auris* aggregates has identified impaired release of daughter cells after cell-division (Borman, Szekely, and Johnson 2016). It was therefore suspected that aggregating cells might exhibit a defect in cell separation due to dysregulation of genes including chitinase (*CTS1*) and chitin synthases (*CHS*) genes (Santana and O’Meara 2021; Short et al. 2019). Our gene expression data showed that a weakly-aggregating isolate (NCPF 13029) expressed more *CTS1* compared to the aggregating strain NCPF 8977. Although *CTS1* expression was higher in the weakly-aggregating strain it is possible this deficiency might be compensated by enhanced expression of chitin (Santana and O’Meara 2021; McCreath, Specht, and Robbins 1995). Some differential levels of *CHS* expression were observed. However, the high potential for post-transcriptional regulation of chitin synthases limits the capacity to make conclusive inferences from these observations. We also did not detect any deletions of core genes (According to Clade I reference genome gene annotation) involved in cell division or separation in sequenced aggregating isolates. Importantly, time lapse microscopy and microfluidics experiments did not reveal obvious defects in cell separation in aggregating and weakly-aggregating strains. Therefore, our data suggests that differences in the aggregation phenotype in our *C. auris* isolates is not governed by defects in cell separation (our hypothesis 1).

A number of biophysical parameters were investigated in this study. Neither cell wall charge nor cell surface hydrophobicity correlated with higher levels of aggregation, despite the importance of both cell wall charge and hydrophobicity to the biology and virulence of fungal pathogens (Gow and Lenardon 2023). We found that aggregation was enhanced at 37°C compared with 30°C and this could explain the finding of aggregates formed in the tissues in a murine model even for poorly aggregating strains (Forgacs *et al*. 2020). The importance of temperature on morphological changes in fungal species including *Candida* and *Histoplasma* is widely acknowledged (Klein and Tebbets 2007; Nguyen and Sil 2008; Shapiro and Cowen 2012). For example, growth at 37°C induces filamentation and biofilm formation in *C. albicans* (Pierantoni et al. 2021; Sudbery 2011). Environmental cues such as increased growth temperature induce a range of bespoke transcription factors and chaperones that govern morphological changes such as hyphal growth and biofilm development (San-Blas et al. 2000; Shapiro et al. 2009; Sudbery, Gow, and Berman 2004). Comprehensive studies on molecular mechanisms that regulate *C. albicans* biofilms have led to the identification of six core transcription factors (Efg1, Tec1, Bcr1, Ndt80, Brg1 and Rob1). Furthermore, 44 additional transcription factors involved in biofilm development have been identified that may be regulated by one or more of the six core biofilm regulators. Two of these core transcription regulators, Efg1 and Tec1 play a crucial role in hyphal induction in *C. albicans*. These studies demonstrate the complexities of transcriptional regulation governing morphological change and complex cellular phenotypes. As with *C. albicans, C. auris* too forms biofilms at 37°C. However, the *C. auris* biofilm primarily comprises yeast cells that produce extracellular matrix (Kean et al. 2018; Sherry et al. 2017). SEM analysis of our aggregating isolates grown at 30°C demonstrated the presence of small amounts of extracellular matrix material that can be associated with the structure of biofilms. We found that the aggregating isolate NCPF 8977 expressed higher level of *XOG1* compared to weakly-aggregating isolate NCPF 13029 at 30°C. Xog1 is an exoglucanase enzyme that has been implicated in formation of extracellular matrix in *C. albicans* (Taff et al. 2012) and may also play a similar role in *C. auris*.

We tested if cell wall amyloid protein(s) are enriched on the surface of strongly-aggregating strains. Cell surface GPI anchored adhesins, like the ALS family of proteins enable adhesion of fungal cells to host surfaces. Our WGS analyses revealed large genomic deletions spanning entire genes (also known as P/A polymorphisms) in weakly-aggregating isolate NCPF 13029 in GPI anchor proteins including Als4, which is a known adhesin in *C. albicans*. Some of these proteins possess the amyloid-forming sequence that promotes protein aggregation (Rauceo et al. 2004; Ramsook et al. 2010). More recently, certain *C. auris* clinical isolates were found to exhibit Als4-mediated aggregation (Bing et al. 2023). We therefore suspected that *C. auris* isolates that aggregate may express more amyloid-proteins than weakly-aggregating isolates. This was confirmed by cell staining with Thioflavin-T, which is an amyloid inhibitor. Prolonged incubation with Thioflavin-T resulted in a dramatic decline in aggregation and adhesion in all isolates. Thioflavin-T did not inhibit aggregation completely and it is therefore likely that aggregation might involve additional factors. For example, the ZAP1 regulon has been shown to contribute to aggregation in *C. albicans* under zinc limitation (Kumar et al. 2017). There may also be a combination of factors that collectively contributed to aggregation that could differ in relative terms in different strains.

In addition to adhesins, yeast cells express Flo proteins and carbohydrate binding lectins on their cell wall that enable aggregation (Govender et al. 2008; Soares 2011; Van Mulders et al. 2009; Touhami et al. 2003). Surface lectins have not yet been identified in the *C. auris* cell wall and it remains to be tested whether carbohydrate-protein interaction may contribute to aggregation. Our HPIC data measuring levels of fundamental cell wall sugars (glucose, mannose and glucosamine) showed comparable carbohydrate profile across all eight isolates and flow cytometric analyses showed strain-specific differences in the exposure of cell wall mannans, chitin and β-glucan – and these did not correlate with the propensity for aggregation. Similarly, sequence analyses of the eight *C. auris* isolates showed strain-specific differences in P/A polymorphisms. RNAseq experiments comparing gene expression in strongly aggregating and weakly-aggregating phenotypes may shed further light on the unidentified surface component(s) that play a role in aggregation (Pelletier 2023).

In this study we have demonstrated that the aggregation phenotype is unlikely to be caused by cell separation defects but may involve cell surface amyloid proteins and other components of the cell wall and extracellular matrix. Our data suggests that aggregation is a complex polygenic property of *C. auris* strains that also responds to key environmental parameters such as environmental temperature.

### Ethical statement

No animals were used in this study that would be covered under Home Office legislation, UK

## Acknowledgements

NG acknowledges support of Wellcome Trust Investigator, Collaborative, Equipment, Strategic and Biomedical Resource awards (101873, 200208, 215599, 224323 and 200208 (the latter awarded to Daan van Aalten)) and the MRC (MR/M026663/2) and the MRC Centre for Medical Mycology (MR/N006364/2) for support. RAF is supported by a Wellcome Trust Career Development Award (225303/Z/22/Z). This study/research is also funded by the National Institute for Health and Care Research (NIHR) Exeter Biomedical Research Centre (BRC). The views expressed are those of the author(s) and not necessarily those of the NIHR or the Department of Health and Social Care.”

## SUPPLEMENTARY FIGURES

Fig. S1: Cellular budding in strongly- and weakly-aggregating isolates. NCPF 8977 (strongly aggregating) and NCPF 8985 (weakly aggregating) were grown in YPD for 24 h at 30°C prior to live cell imaging in YPD containing 1 µg/ml Calcoflour White (CFW) for 8 h. Representative movies demonstrate comparable cellular budding and cell separation in both isolates. Movie S1 A) NCPF 8977 in CFW channel. Movie S1 B) NCPF 8985 in CFW channel. Scale bar represents 5 µm.

Fig. S2: Gating strategies. A) Gating strategy for adhesion assay. Gate P1 represents uncoated carboxylated yellow-green fluorescent polystyrene microspheres only, Gate P2 represents *C. auris* cells without polystyrene microspheres. Representative density plot (far right) shows systematic identification of *C. auris* cells that have adhered to polystyrene microspheres. B) Gating strategy for cell wall staining. Representative density plots showing whole population (left) of *C. auris* cells using side scatter area on y-axis and forward scatter area on x-axis. Single-cells population (right) identified using forward scatter height on y-axis and forward scatter area on x-axis.

