## Supplementary figures and images for "Strain and temperature dependent aggregation of *Candida auris* is attenuated by inhibition of surface amyloid proteins"

### Supplementary Figure 2

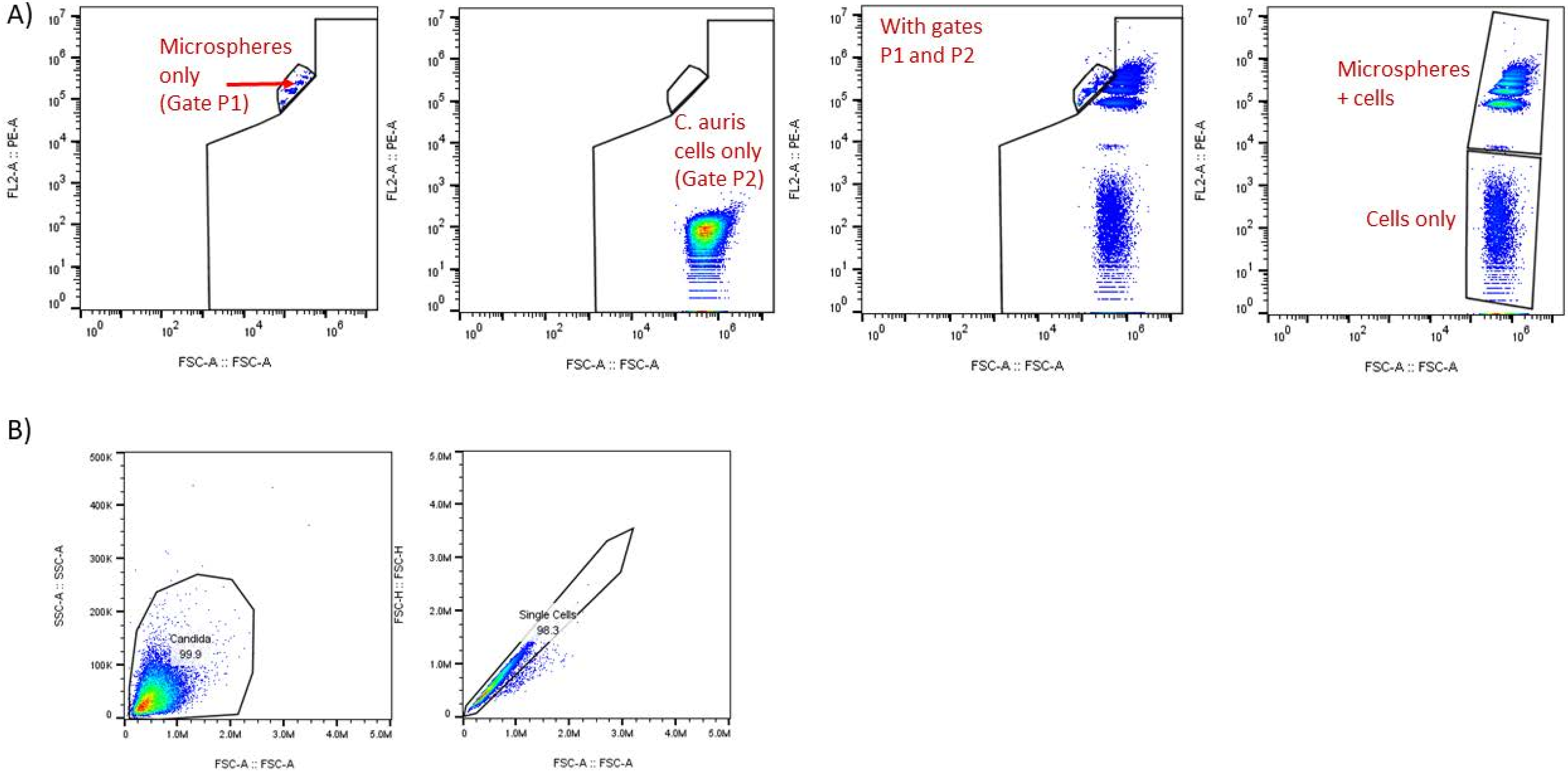
